# The mycobacterial cell wall partitions the plasma membrane to organize its own synthesis

**DOI:** 10.1101/544338

**Authors:** Alam García-Heredia, Takehiro Kado, Caralyn E. Sein, Julia Puffal, Sarah H. Osman, Julius Judd, Todd A. Gray, Yasu S. Morita, M. Sloan Siegrist

## Abstract

Many antibiotics target the assembly of cell wall peptidoglycan, an essential, heteropolymeric mesh that encases most bacteria. Different species have characteristic subcellular sites of peptidoglycan synthesis that they must carefully maintain for surface integrity and, ultimately, viability. In rod-shaped bacteria, cell wall elongation is spatially precise yet relies on a limited pool of lipid-linked precursors that generate and are attracted to membrane disorder. By tracking enzymes, substrates and products of peptidoglycan biosynthesis in *Mycobacterium smegmatis*, we show that precursors are made in plasma membrane domains that are laterally and biochemically distinct from sites of cell wall assembly. Membrane partitioning is required for robust, orderly peptidoglycan synthesis, indicating that these domains help template peptidoglycan synthesis. The cell wall-organizing protein DivIVA and the cell wall itself are essential for domain homeostasis. Thus, the peptidoglycan polymer feeds back on its membrane template to maintain an environment conducive to directional synthesis. We further show that our findings are applicable to rod-shaped bacteria that are phylogenetically distant from *M. smegmatis*, demonstrating that horizontal compartmentalization of precursors is a general feature of bacillary cell wall biogenesis.

## Main

The final lipid-linked precursor for peptidoglycan synthesis, lipid II, is made by the glycosyltransferase MurG in the inner leaflet of the plasma membrane. Lipid II is then flipped to the outer leaflet by MurJ where its disaccharide-pentapeptide cargo is inserted into the existing cell wall by membrane-bound transglycosylases and transpeptidases (Zhao et al., 2017). Early *in vitro* work in *Staphylococcus aureus* and *Escherichia coli* indicated that a fluid microenvironment might stimulate the activities of MurG and the upstream, lipid I synthase MraY (Norris & Manners, 1993). More recent *in vivo* data has localized *Bacillus subtilis* MurG to **r**egions of **i**ncreased **f**luidity (**RIF**s, [Muller et al., 2016; Strahl et al., 2014]), one of three classes of membrane domains that have been described in bacteria to date. In mycobacteria, **i**ntracellular **m**embrane **d**omains (IMD, formerly called PMf, [Morita et al., 2005]) can be separated from the conventional plasma membrane (PM-CW, for **p**lasma **m**embrane associated with **c**ell **w**all) by sucrose density gradient fractionation. The proteome and lipidome of IMD are distinct from those of the PM-CW (Hayashi et al., 2016; Morita et al., 2005). Reanalysis of our proteomics data (Hayashi et al., 2016) suggested that *Mycobacterium smegmatis* MurG is enriched in the IMD while sequentially-acting transglycosylases and transpeptidases associate with the PM-CW. While PM-CW-resident proteins distribute thorough the perimeter of live mycobacteria, IMD-resident proteins localize along the sidewall but are enriched toward sites of polar cell elongation (Hayashi et al., 2016; Hayashi et al., 2018). We also noted that polar enrichment of MurG resembles that of the validated IMD marker GlfT2 (Hayashi et al., 2016; Meniche et al., 2014), but that nascent peptidoglycan at the mycobacterial poles primarily abuts rather than colocalizes with GlfT2 (Hayashi et al., 2018). These observations suggested a model where lipid II synthesis is segregated from subsequent steps of cell wall assembly (Fig. 1A).

**Fig. 1.**
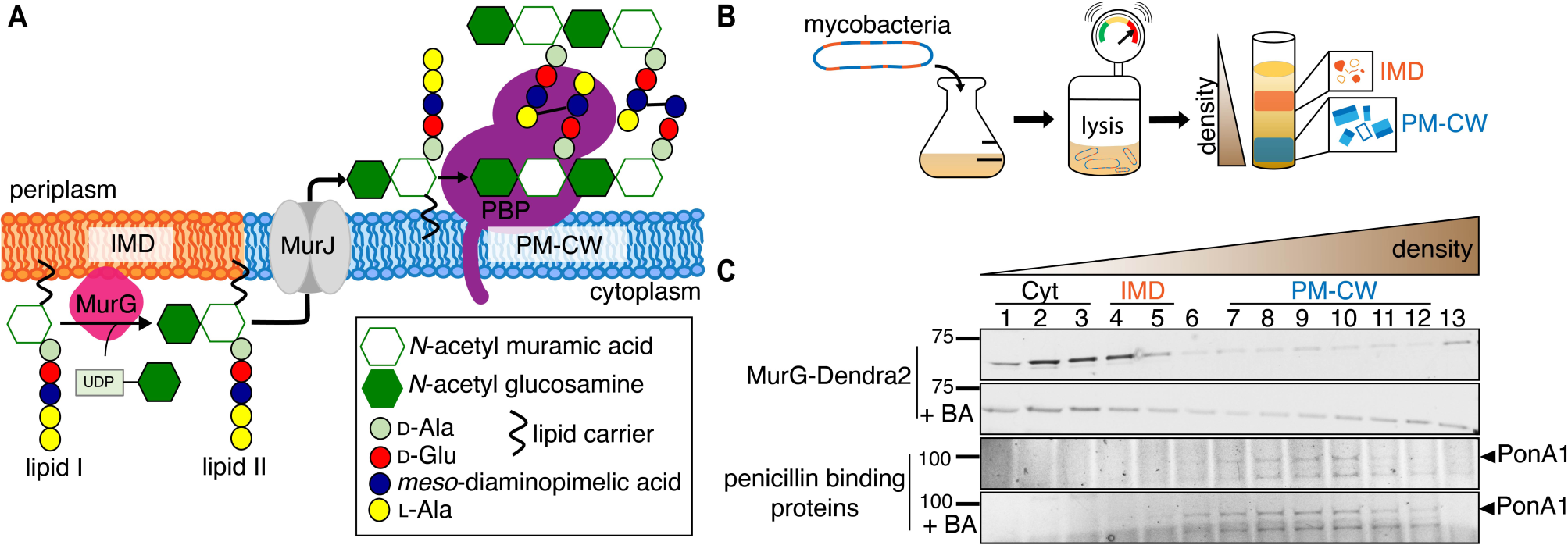
MurG is enriched in the IMD and PBPs associate with PM-CW. (A) Membrane-bound steps of peptidoglycan synthesis with hypothesized partitioning into IMD and PM-CW. (B) Bacteria are lysed by nitrogen cavitation and cell lysate is sedimented on a sucrose density gradient. (C) Lysates from wild-type or MurG-Dendra2-expressing *M. smegmatis* were fractionated as in (B) and separated by SDS-PAGE. Top, in-gel fluorescence shows MurG-Dendra2 association with the IMD. Treatment with benzyl alcohol (BA) redistributed the protein across the fractions. Bottom, wild-type *M. smegmatis* membrane fractions were incubated with Bocillin-FL prior to SDS-PAGE. Labeled PBPs are enriched in PM-CW.

To test this model, we first expressed a functional MurG-Dendra2 fusion in *M. smegmatis* (Figure 1–figure supplement 1A) and assayed its distribution in membrane fractions that had been separated by density gradient (Fig. 1B). MurG-Dendra2, a peripheral membrane protein, was enriched in both the cytoplasmic and IMD membrane fractions (Fig. 1C, top; Figure 1 – figure supplement 2A). In intact cells, polar enrichment of MurG-Dendra2 was coincident with that of the IMD marker mCherry-GlfT2 (Figure 2–figure supplement 1). This spatial relationship was similar to that previously observed for MurG and GlfT2 (Hayashi et al., 2016; Hayashi et al., 2018; Meniche et al., 2014).

**Fig. 2.**
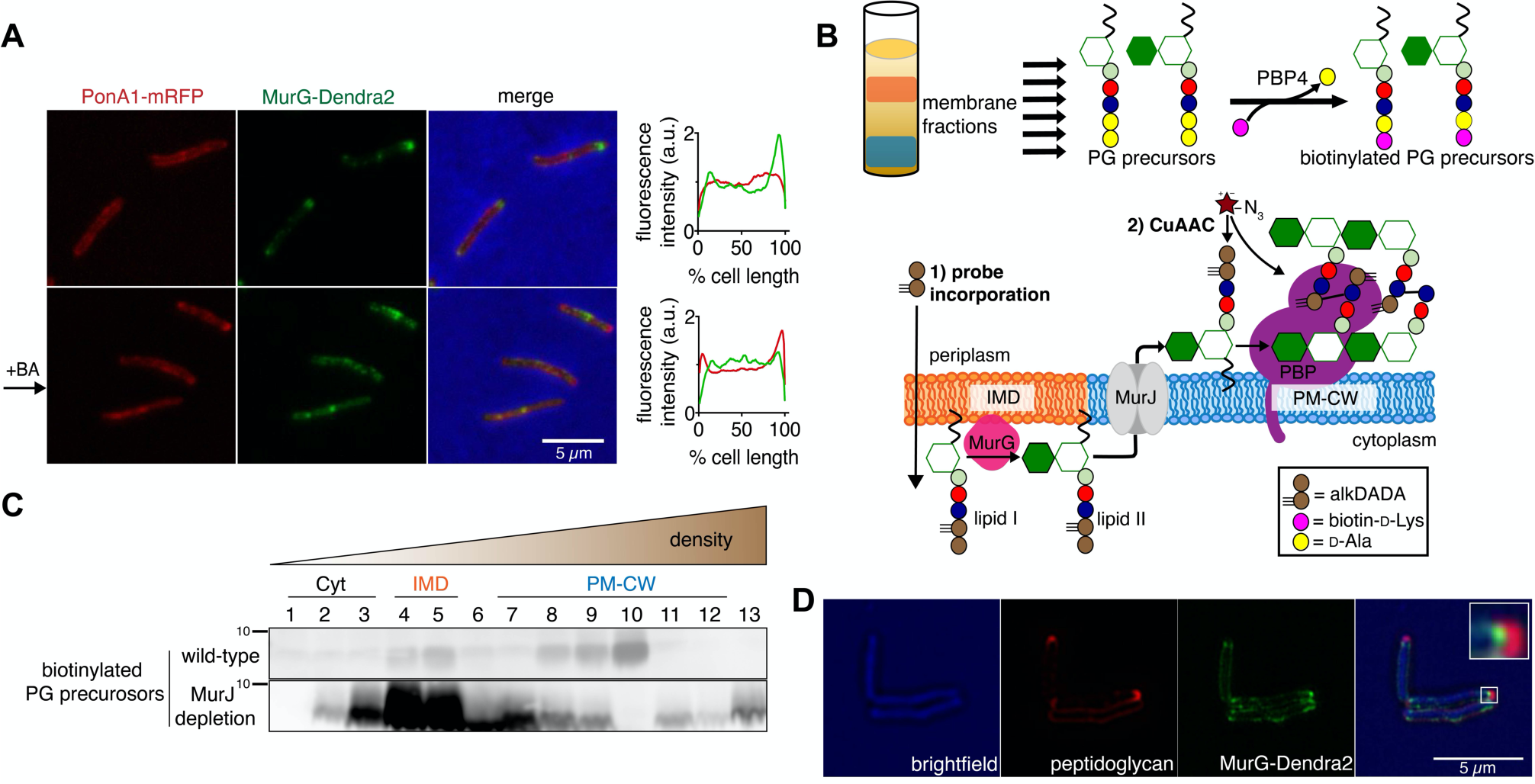
Lipid II is synthesized in the IMD and trafficked to the PM-CW. (A) Left, conventional microscopy of *M. smegmatis* coexpressing PonA1-mRFP and MurG-Dendra2 treated +/- benzyl alcohol (BA). Right, fluorescence distribution of the fusion proteins. a.u., arbitrary units. (B) Top, detection scheme for lipid-linked peptidoglycan (PG) precursors from membrane fractions. Bottom, metabolic labeling scheme of mycobacterial cell wall synthesis (García-Heredia et al., 2018). (C) PG precursors are labeled as in (B), top. The labeled precursors are in the IMD and PM-CW of wild-type *M. smegmatis* but accumulate in the IMD upon MurJ depletion (García-Heredia et al., 2018). (D) *M. smegmatis* expressing MurG-Dendra2 were incubated with alkDADA. Surface-exposed alkynes on fixed cells were detected by CuAAC (García-Heredia et al., 2018). Cells imaged by structured illumination microscopy (SIM-E).

Enzymes from the **p**enicillin **b**inding **p**roteins (PBPs) and **s**hape, **e**longation, **d**ivision and **s**porulation (SEDS) families integrate the disaccharide-pentapeptide from lipid II into peptidoglycan (Zhao et al., 2017). While our proteomics did not detect many polytopic membrane proteins, including SEDS proteins, our PM-CW dataset was enriched for all known PBPs (Hayashi et al., 2016). Fluorescent derivatives of β-lactam antibiotics, such as Bocillin-FL, bind to PBPs and report transpeptidase-active enzymes. We incubated subcellular fractions from wild-type *M. smegmatis* with Bocillin-FL and identified fluorescent proteins in the PM-CW (Fig. 1C, bottom; Figure 1 –figure supplement 2B). As expected for PBPs, the signal from these bands was diminished by pre-treatment with ampicillin (Figure 1–figure supplement 3). We focused on characterizing PonA1, an essential bifunctional transglycosylase/transpeptidase in *M. smegmatis* (Hett et al., 2010; Kieser et al., 2015). Depletion of PonA1 (Hett et al., 2010) resulted in the loss of the higher molecular band (Figure 1–figure supplement 3), confirming this protein is present and active in the PM-CW (Fig. 1C, bottom). We next expressed a functional PonA1-mRFP fusion in *M. smegmatis* (Kieser et al., 2015) and found that it was distributed around the cell perimeter in a manner similar to the functional PM-CW marker PimE-GFP (Figure 1–figure supplement 1B; Figure 2–figure supplement 1, [Hayashi et al., 2016]). Coexpression of MurG-Dendra2 and PonA1-mRFP confirmed that the proteins are laterally segregated from one another (Fig 2A). Together, our data show that MurG and PonA1 occupy membrane compartments that are biochemically and spatially distinct.

The association of MurG with the IMD and of PonA1 with the PM-CW implied that the IMD is the site of lipid II synthesis while the PM-CW is where peptidoglycan assembly takes place. We refined an *in vitro* D-amino acid exchange assay to detect lipid-linked peptidoglycan precursors from membrane fractions (Fig. 2B, [García-Heredia et al., 2018; Qiao et al., 2014]). In wild-type cells, we detected biotinylated molecules in both the IMD and PM-CW (Fig. 2C; Figure 1–figure supplement 2B). We hypothesized that the labeled species comprise precursors in both the inner and outer leaflet of the plasma membrane. We and others have shown that depletion of MurJ results in accumulation of biotinylated precursors (García-Heredia et al., 2018; Qiao et al., 2017). By performing the D-amino acid exchange reaction on membrane fractions obtained from MurJ-depleted *M. smegmatis* (Figure 1–figure supplement 1C), we found that precursors accumulate in the IMD (Fig. 2C; Figure 1–figure supplement 2C). These results suggest that lipid II is made in the IMD and transferred to the PM-CW in a MurJ-dependent manner.

Based on our biochemical data, we hypothesized that lipid II incorporation into the cell wall is laterally segregated from its synthesis. We previously showed that alkynyl and azido D-amino acid dipeptides (Liechti et al., 2014) incorporate into lipid-linked peptidoglycan precursors in *M. smegmatis* (García-Heredia et al., 2018) and that metabolic labeling with alkynyl dipeptide (alkDADA or EDA-DA, [Liechti et al., 2014]) is most intense in regions adjacent to the IMD marker mCherry-GlfT2 (Hayashi et al., 2018). We labeled MurG-Dendra2-expressing *M. smegmatis* with alkDADA and detected the presence of the alkyne by copper-catalyzed azide-alkyne cycloaddition (CuAAC, [García-Heredia et al., 2018]). To distinguish extracellular alkynes present in periplasmic lipid II and newly-polymerized cell wall from alkynes originating from cytoplasmic lipid II, we selected picolyl azide-Cy3 as our label because of its poor membrane permeability (Fig. 2B, [Yang & Hinner, 2015]). Using this optimized protocol, we observed nascent peptidoglycan deposition at the polar tip, whereas MurG-Dendra2 was proximal to this site (Fig. 2D). Our data suggest that lipid II synthesis is laterally partitioned from the subsequent steps of peptidoglycan assembly. MurJ depletion reduced and delocalized alkDADA-derived fluorescence (Figure 2–figure supplement 2), consistent with a gatekeeper role for the flippase in both lateral membrane compartmentalization and flipping across the inner membrane.

We next wanted to understand the effect of partitioning on cell wall synthesis. We opted to perturb the plasma membrane with benzyl alcohol, which has been used to disperse membrane domains in animal and bacterial cells, including *B. subtilis* RIFs (Muller et al., 2016; Nagy et al., 2007). In *M. smegmatis*, we found that benzyl alcohol reduced the cellular material associated with the IMD (Fig 3A, Figure 1–figure supplement 2D) and altered the distribution of FM4-64, a non-specific lipophilic dye (Figure 3–figure supplement 1A) and of plasma membrane glycolipids (Figure 3–figure supplement 2). However, benzyl alcohol did not alter labeling by N-AlkTMM or O-AlkTMM (Figure 3–figure supplement 1B), probes that respectively mark the noncovalent and covalent lipids of the outer ‘myco’ membrane (Foley et al., 2016). These observations suggest that benzyl alcohol primarily affects the plasma membrane. MurG-Dendra2 was also notably less enriched in the IMD fraction following benzyl alcohol treatment (Fig. 1C, top; Figure 1–figure supplement 2E) and, in live cells, at the poles (Fig. 2A). By contrast, benzyl alcohol produced subtle changes in the subcellular distribution of active PBPs (Fig. 1C, bottom; Figure 1–figure supplement 2D), although PonA1 shifted toward the poles in live cells (Fig. 2A). Disruption of plasma membrane architecture was accompanied by dampening and delocalization of peptidoglycan assembly (Fig. 3B) as well as a reduction in lipid precursor synthesis (Fig. 3C) and halt in polar elongation (Figure 3–figure supplement 1B). The effects of benzyl alcohol were reversible, as indicated by colony forming units and prompt recovery of peptidoglycan synthesis (Figure 3–figure supplement 3). Dibucaine, a membrane-fluidizing anesthetic (Papahadjopoulos et al., 1975), also delocalized IMD-resident proteins (Figure 3–figure supplement 4) and reduced peptidoglycan synthesis (Fig. 3B). These data suggest that membrane architecture is essential feature of both peptidoglycan synthesis and cell growth in *M. smegmatis*.

**Fig. 3.**
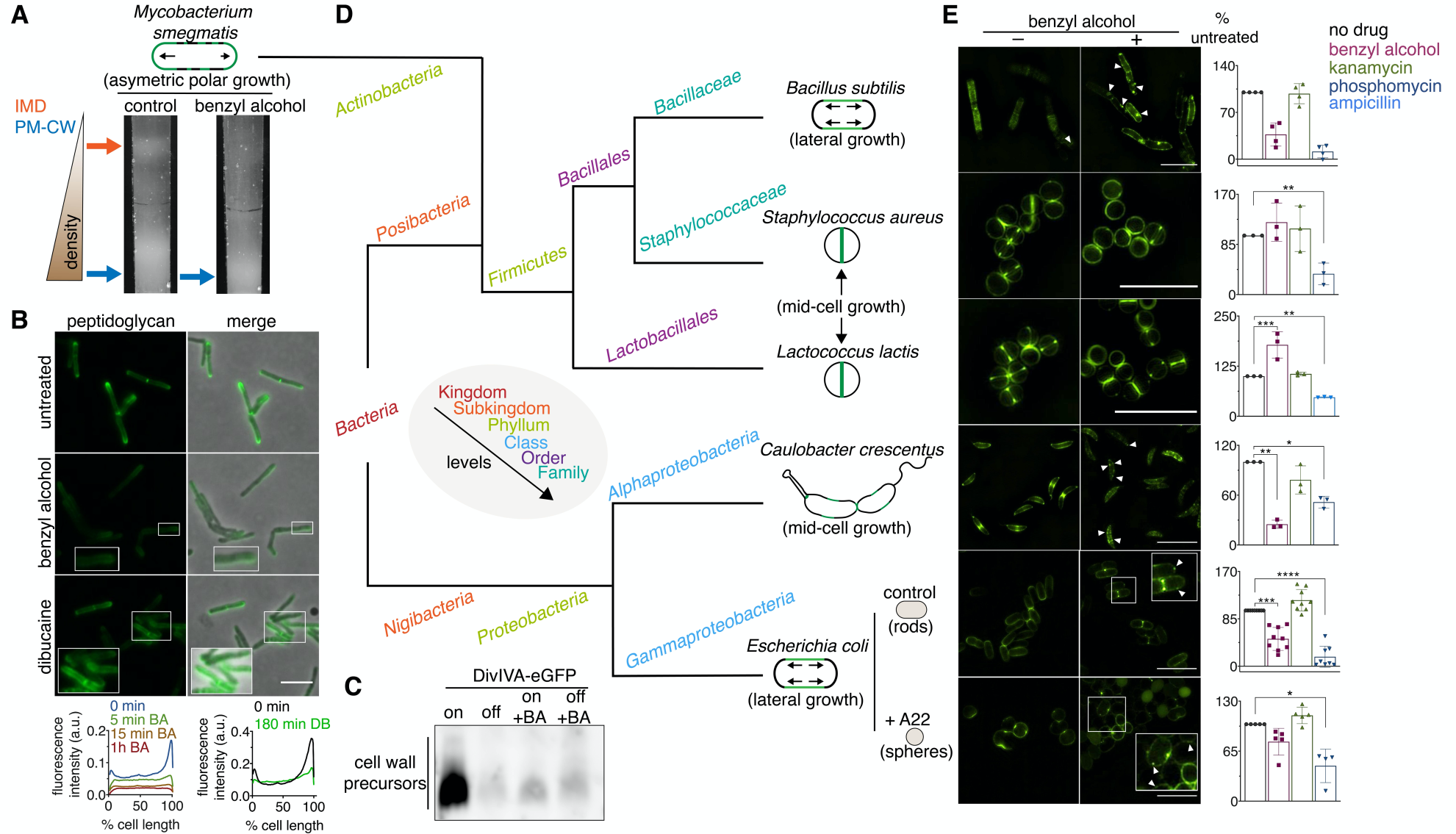
Membrane perturbations disrupt peptidoglycan biogenesis in *M. smegmatis* and phylogenetically-distant bacilli. (A) Lysates from wild-type *M. smegmatis* treated +/- benzyl alcohol (BA) were sedimented in a sucrose density gradient. (B) Top, wild-type *M. smegmatis* was incubated or not with benzyl alcohol or dibucaine, then labeled with alkDADA. Bottom, the distribution of peptidoglycan labeling from wild-type *M. smegmatis* that was incubated with BA or dibucaine (DB) for the indicated time was quantitated as in Fig. 2A, except that signal intensity was not normalized. (C) DivIVA-eGFP-ID *M. smegmatis* was either treated with benzyl alcohol, depleted of DivIVA, or both, and the peptidoglycan precursors from whole cells were biotinylated as in Fig. 2B, top. (D) Phylogenetic tree constructed with 16S rDNA sequences (rate of mutation not considered). Taxonomic groups matched with colors to their levels with only diverging points shown. Shapes and growth modes illustrated for select species. (E) Left, different bacteria treated +/- benzyl alcohol followed by alkDADA incubation. Arrowheads highlight irregular patches of peptidoglycan. Insets are magnified. Where applicable, *E. coli* was preincubated with A22. Right, bacteria were treated with benzyl alcohol, translation-inhibiting kanamycin, or peptidoglycan-acting phosphomycin or ampicillin then labeled as in (B) and analyzed by flow cytometry (see Methods). Median fluorescence intensity (MFI) values were normalized to untreated controls and experiments were performed 3-9 times in triplicate. Error bars, +/- standard deviation of biological replicates. *, p < 0.05; **, p < 0.005; ***, p < 0.0005; ****, p < 0.00005, ratio paired t-tests and one-way ANOVA with Dunnet’s test for non-normalized MFI of biological replicates. Scale bars, 5 µm.

Benzyl alcohol delocalized or dampened cell wall assembly in other rod-shaped bacteria with divergent envelope composition and modes of growth (Fig. 3D-E). Peptidoglycan synthesis was less affected by benzyl alcohol in coccoid species, which lack MreB, and in rounded, A22-treated *E. coli*, which have MreB inhibited (Fig. 3E). Thus, membrane partitioning is necessary for effective, directional cell wall synthesis in rod-shaped bacteria.

How is the IMD partitioned away from the rest of the plasma membrane? The architecture of eukaryotic membranes is influenced by transient links, or pinning, to the cytoskeleton (Fujimoto & Parmryd, 2016; Liu et al., 2015). In *B. subtilis* and *E. coli*, actin homologs like MreB organize RIFs (Oswald et al., 2016; Strahl et al., 2014) and direct peptidoglycan synthesis along the lateral cell surface (Daniel & Errington, 2003; Iwai et al., 2002; Shi et al., 2018; Zhao et al., 2017). While MreB is absent in mycobacteria, the essential tropomyosin-like protein DivIVA concentrates cell wall assembly at the poles (Daniel & Errington, 2003; Iwai et al., 2002; Melzer et al., 2018; Shi et al., 2018; Zhao et al., 2017). DivIVA depletion results in deformation and rounding of mycobacterial cells (Kang et al., 2008). Given the similarities in DivIVA and MreB function, we hypothesized that DivIVA creates and/or maintains the IMD. We used *M. smegmatis* expressing DivIVA-eGFP-ID (in which DivIVA is fused to both eGFP and an inducible degradation tag [Meniche et al., 2014]) to deplete DivIVA. Depletion of the protein reduced the amount of IMD-associated cellular material (Fig. 4A, Figure 1–figure supplement 2F-G), altered the distribution of plasma membrane glycolipids (Figure 3–figure supplement 2), and delocalized the IMD marker mCherry-GlfT2 from the poles (Fig. 4B). DivIVA phosphorylation regulates MraY and/or MurG activity via an indirect, unknown mechanism (Jani et al., 2010). Consistent with these data, we found that depletion of the protein reduced lipid-linked peptidoglycan precursor abundance (Fig. 3C). The suppressive effects of benzyl alcohol and DivIVA depletion on precursor abundance were not additive (Fig. 3C), suggesting that the perturbations act on the same pathway.

**Fig. 4.**
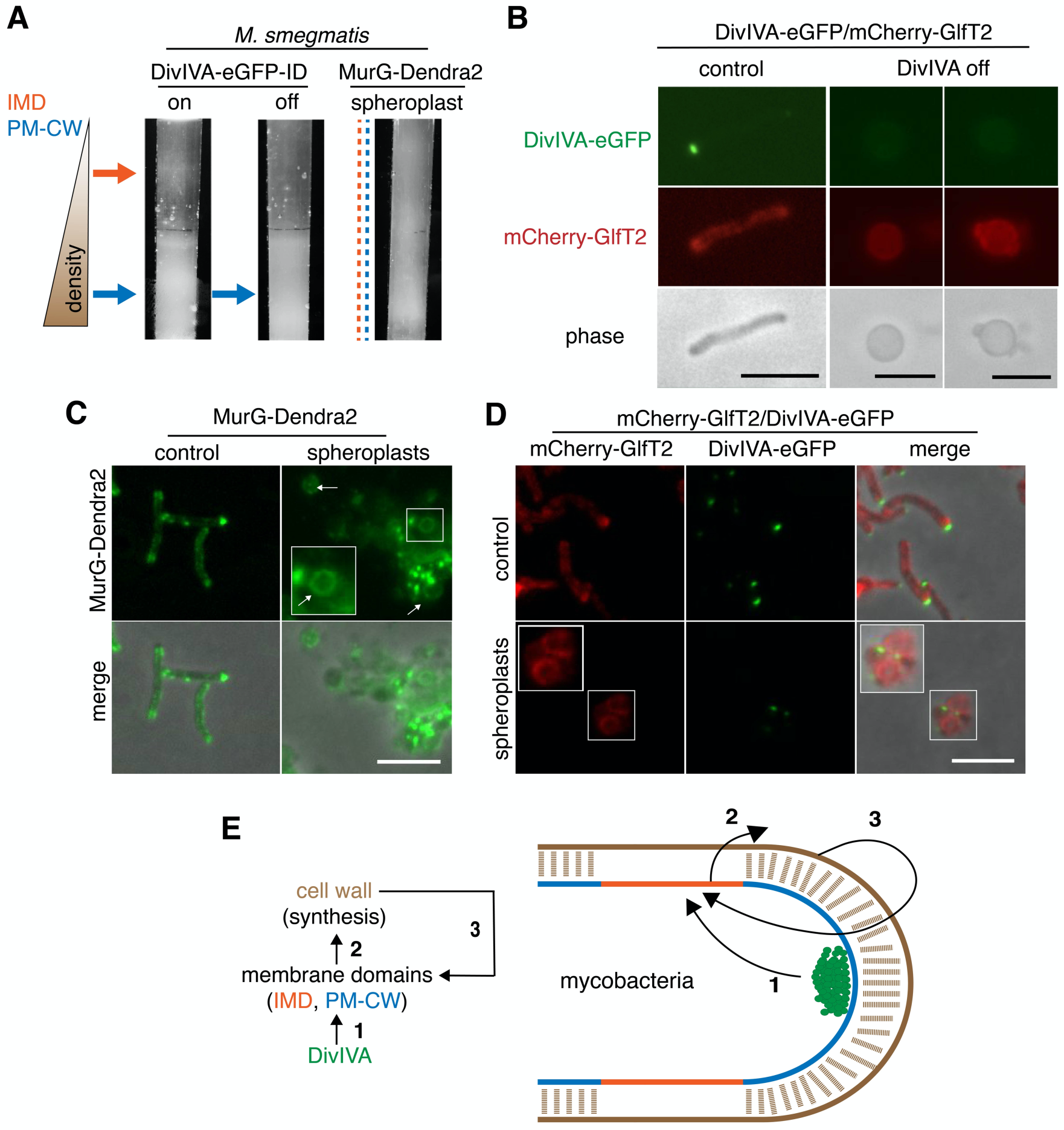
Plasma membrane domains are maintained by DivIVA and the cell wall. (A) Lysates from MurG-Dendra2- or DivIVA-eGFP-ID-expressing *M. smegmatis* that had been depleted of DivIVA (Meniche et al., 2014) or spheroplasted (Melzer et al., 2018), were sedimented in a density gradient. (B). DivIVA was depleted or not in *M. smegmatis* expressing both DivIVA-eGFP-ID and mCherry-GlfT2. Normally, DivIVA localizes at the polar tip and mCherry-GlfT2 concentrates proximal to DivIVA. Depletion of DivIVA delocalizes mCherry-GlfT2. *M. smegmatis* expressing MurG-Dendra2 (C) or coexpressing mCherry-GlfT2 and DivIVA-eGFP-ID (D) were spheroplasted or not and imaged. In spheroplasted cells, the IMD-associated proteins distribute along the cell periphery. Arrows mark spheroplasts outside and within insets, which have increased size and brightness. (E) Model for self-organization of plasma membrane and cell wall in *M. smegmatis*. Brown line indicates the cell wall. Short brown lines perpendicular to the membrane and cell wall indicate that the cell wall is likely to be physically connected to the membrane in the PM-CW regions (Morita et al., 2005). All scale bars, 5 µm.

As lipid II both generates and homes to disordered regions of model membranes (Ganchev et al., 2006; Jia et al., 2011; Valtersson et al., 1985), the effect of DivIVA on precursors suggests that concentrated peptidoglycan synthesis is a cause or a consequence (or both) of IMD/PM-CW partitioning. In other organisms, lipid II production is required for MreB rotation (Dominguez-Escobar et al., 2011; Garner et al., 2011; van Teeffelen et al., 2011) and to recruit MreB to the plasma membrane (Schirner et al., 2015), so the precursor might also play an indirect role in compartmentalizing the mycobacterial membrane via its influence on DivIVA. However, we previously found that the IMD is biochemically intact after 8 hours of treatment with D-cycloserine (Hayashi et al., 2018), an antibiotic that we have shown to block *M. smegmatis* peptidoglycan precursor synthesis within 1 hour (García-Heredia et al., 2018). IMD-resident proteins delocalize, but not until 6 hours of treatment. The persistence of IMD-resident proteins and the time frame of delocalization indicate that concentrated lipid II synthesis is more likely a consequence, rather than a cause, of mycobacterial membrane compartmentalization.

Our data supported a model in which DivIVA is necessary to maintain membrane partitioning, which in turn supports efficient synthesis of peptidoglycan precursors and their precise incorporation into the cell wall. However, we noted that eukaryotic membrane architecture can also be influenced by transient interactions with external structures like extracellular matrix and cellulose (Fujimoto & Parmryd, 2016; Liu et al., 2015). Furthermore, biophysical modeling of bacteria suggests that osmotic pinning of the plasma membrane against the cell wall can induce microphase separation (Mukhopadhyay et al., 2008). In mycobacteria, co-fractionation of the **p**lasma **m**embrane and **c**ell **w**all (*i.e.* PM-CW) upon mechanical cell lysis implies that they are physically connected (Morita et al., 2005). We wondered whether the peptidoglycan polymer itself might contribute to the membrane partitioning that organizes its synthesis. To test this hypothesis, we spheroplasted (enzymatically removed the cell wall; Melzer et al., 2018) *M. smegmatis* that expressed MurG-Dendra2 or DivIVA-eGFP-ID and mCherry-GlfT2. All fusions were functional (Figure 1–figure supplement 1, [Hayashi et al., 2016; Meniche et al., 2014]). Fractionated lysates from spheroplasted mycobacteria had more diffuse distribution of cellular material and indistinct separation of the IMD and PM-CW fractions (Fig. 4A). Consistent with the macroscopic appearance of the fractionated lysate, MurG-Dendra2 was distributed throughout the gradient (Figure 4–figure supplement 1). MurG-Dendra2 and the IMD marker mCherry-GlfT2 were also diffusely distributed around the periphery of spheroplasted cells while DivIVA, a non-IMD protein, remained in foci (Fig. 4C-D). These experiments indicate that an intact cell wall and DivIVA are both required for membrane compartmentalization in *M. smegmatis* (Fig. 4E).

While peptidoglycan biogenesis is well known to vertically span the inner and outer leaflets of the plasma membrane, here we demonstrate in *M. smegmatis* that it is also horizontally partitioned. Partitioning of the mycobacterial membrane by DivIVA and the cell wall follows similar logic to that of eukaryotic membranes, which can be compartmentalized by pinning to cytoplasmic structures such as the actin cytoskeleton and to external structures like extracellular matrix and cellulose (Fujimoto & Parmryd, 2016; Liu et al., 2015), and of model lipid bilayers, which can be phase separated by adhesive forces (Gordon et al., 2008; Mukhopadhyay et al., 2008). In mycobacteria, however, the membrane regions that promote cell wall synthesis are segregated by the end product of the pathway (Fig. 4E). For these and other rod-shaped bacteria, our model is that the membrane-cell wall axis is a self-organizing system in which directed cell wall synthesis organizes the plasma membrane, and an organized plasma membrane in turn makes cell wall elongation more efficient and precise.

## Acknowledgments

We are grateful to Drs. James Chambers and Amy Burnside for microscopy and flow cytometry guidance, Drs. Eric Strieter and Jiaan Liu and Ms. Sylvia Rivera and Katherine Chacón-Vargas for technical assistance. We also thank Drs. Hesper Rego and Eric Rubin for *ponA1-mRFP M. smegmatis*, Dr. Christopher Sassetti for the MurJ and MurG depletion strains, and Dr. Benjamin Swarts for O-AlkTMM and N-AlkTMM. Research was supported by funds from the National Institutes of Health (NIH) under awards R21 AI144748 (Y.S.M and M.S.S.), U01 CA221230 and NIH DP2 AI138238 (M.S.S.) and R03 AI140259-01 (Y.S.M.), and R01 AI097191 (J.J. and T.A.G.). A.G.-H. was supported by an Honors Fellowship from Universidad Autónoma de Nuevo León. J.P. is a recipient of the Science Without Boarders Fellowship from CAPES-Brazil (0328-13-8). T.K. was supported by a postdoctoral fellowship from Uehara Memorial Foundation.

## Contributions

M.S.S., A.G.-H. and Y.S.M. designed the experiments and analyzed the data. A.G.-H. performed most of the experiments. T.K. constructed the DivIVA-eGFP-ID/mCherry-GlfT2 *M. smegmatis* mutant and helped with experiments from Figs. 3B, 4B and Figure 3–figure supplement 4. J.P. performed immunoblots for Figure 1–figure supplement 2 and thin-layer chromatography for Figure 3–figure supplement 2. C.E.S performed experiments for Fig. 4C. S.H.O. did experiments for Fig. 4A and Figure 4–figure supplement 1. Y.S.M. did the glycolipid analysis for Figure 1–figure supplement 1. J.J. and T.A.G. constructed the MurG-Dendra2 plasmid. M.S.S. and A.G.-H. wrote the manuscript. All authors reviewed and edited the manuscript.

## Data and materials availability

All data are available in the main text or the supplementary materials. Raw data is available at open science framework through the next link: https://osf.io/fm794/?view_only=7d5a24b82dd94744a46420aee09bad7f

Figure supplements

Supplementary information: Tables S1-4

## Methods

### Bacterial strains and growth conditions

*Mycobacterium smegmatis* mc^2^155 was grown in Middlebrook 7H9 growth medium (BD Difco, Franklin Lakes, NJ USA) supplemented with 0.4% (v/v) glycerol, 0.05% (v/v) Tween-80 (Sigma-Aldrich, St. Louis, MO USA) and 10% ADC (albumin-dextrose-catalase) as well as apramycin (50 µg/mL), kanamycin (25 μg/mL; Sigma-Aldrich, St. Louis, MO USA) and hygromycin (50 μg/mL) where appropriate. *Staphylococcus aureus* ATCC BA-1718 was grown in BHI (BD Difco, Franklyn Lakes NJ, USA); *Escherichia coli* K12 and *Bacillus subtilis* ZB307, in LB (VWR, Radnor PA, USA); *Caulobacter crescentus* NA 1000, in peptone yeast extract (BD Difco, Franklyn Lakes NJ, USA); and *Lactococcus lactis* NRRL B633, in MRS (Oxoid, Basingstoke, Hampshire UK). All bacteria were grown shaking at 37°C with the exception of *C. crescentus*, which was incubated at 30°C. See Table S3 for more information.

### Mutant strain construction

To test the function of PimE-GFP-FLAG fusion, an expression vector for PimE-GFP-FLAG (pYAB186) (Hayashi et al., 2016) was electroporated into a *pimE* deletion mutant (Morita et al., 2006). Three independent colonies were picked for the phenotypic complementation of AcPIM6 biosynthetic defects. Lipid purification and analysis were performed as described previously (Morita et al., 2006).

The *murG* gene was amplified by PCR from *M. smegmatis* mc^2^155 genomic DNA by PCR, excluding the stop codon, and was inserted into pMSR in-frame with mycobacterial codon-optimized Dendra2 using In-Fusion cloning (Takara Bio, Mountain View, CA USA). This construct was transformed by electroporation into *M. smegmatis* mc^2^155, where it integrates at the L5 *attB* site, and was selected by apramycin treatment. Constitutive expression of the MurG-Dendra2 fusion was achieved through the Psmyc promoter (GenBank: AF395207.1). The plasmid construct was validated by Sanger sequencing.

To replace the endogenous *glfT2* gene with a gene encoding HA-mCherry-GlfT2 in the DivIVA-eGFP-ID strain, we electroporated pMUM052 (Hayashi et al., 2016) into DivIVA-eGFP-ID *M. smegmatis*, and positive clones were isolated using hygromycin resistance marker and SacB-dependent sucrose sensitivity. Correct replacement of the *glfT2* gene was confirmed by PCR.

### Generation of spheroplasts

To generate spheroplasts, we followed a previous protocol (Melzer et al., 2018). Briefly, wild-type, MurG-Dendra2-expressing or mCherry-GlfT2/DivIVA-eGFP-ID-coexpressing *M. smegmatis* was grown until log phase. Glycine (1.2% w/v final concentration) was added and the culture was incubated for 24 h at 37°C with shaking. Afterwards, the cells were washed with sucrose-MgCl_2_-maleic acid (SMM) buffer (pH 6.8) and harvested by centrifugation (4000 x *g* for 5 min). The pellet was resuspended in 7H9 medium where the water was replaced with SMM buffer; the medium also was supplemented with glycine (1.2% final concentration) and lysozyme (50 µg/mL final concentration). Bacteria were incubated another 24 h at 37°C with shaking, and then spheroplasts where either imaged by conventional fluorescence microscopy or lysed by nitrogen cavitation immediately for subsequent biochemical analysis.

### Membrane fractionation

Log phase *M. smegmatis* (that, where applicable, were untreated, treated with benzyl alcohol, depleted for DivIVA or spheroplasted; see Table S1) were harvested by centrifugation and washed in PBS + 0.05% Tween-80 (PBST). One gram of wet pellet was resuspended in 5 mL of lysis buffer containing 25 mM HEPES (pH 7.4), 20% (w/v) sucrose, 2 mM EGTA, and a protease inhibitor cocktail (Thermo Fisher Scientific, Waltham, MA USA) as described (Morita et al., 2005). Bacteria were lysed using nitrogen cavitation at 2000 psi for 30 min three times. Cell lysates were centrifuged at 3220 x *g* for 10 min at 4°C twice to remove unlysed cells prior to loading on a 20-50% sucrose gradient. Membrane-containing fractions were collected in 1 mL after ultracentrifugation at 35,000 rpm on an SW-40 rotor (Beckman, Brea, CA USA) for 6 hours at 4°C and stored at -80°C prior to analysis.

### Detection of proteins in membrane fractions

MurG-Dendra2 and penicillin-binding proteins (PBPs) were detected by in-gel fluorescence. For MurG-Dendra2, membrane fractions were incubated with an equal volume of 2x loading buffer and then separated by SDS-PAGE on a 12% polyacrylamide gel. To detect PBPs, 50 µg total protein from wild-type *M. smegmatis* membrane fractions were incubated with 40 µM of Bocillin-FL (Thermo Fisher Scientific, Waltham, MA USA) for 30 min in the dark at 37°C. An equal volume of 2x loading buffer was then added and the mixture was boiled for 3 min at 95°C and then incubated on ice for 30 min. Membrane mixtures were separated on a 12% polyacrylamide gel. Gels were washed in distilled water and imaged using an ImageQuant LAS 4000mini (GE Healthcare, Chicago, IL USA).

MurJ, PimB’ and MptA were detected by immunoblot. Briefly, cell lysate or membrane fraction proteins were separated by SDS-PAGE on a 12% polyacrylamide gel and transferred to a PVDF membrane. The membrane was blocked with 3% milk in PBS + 0.05% Tween-80 (PBST) and then incubated overnight with primary antibodies (monoclonal mouse anti-FLAG [to detect MurJ; Sigma], and polyclonal rabbit anti-PimB’ or anti-MptA antibodies (Sena et al., 2010). Antibodies were detected with appropriate secondary antibodies conjugated to horseradish peroxidase (GE Healthcare, Chicago, Il USA). Membranes were rinsed in PBS + 0.05% Tween-20 and visualized by ECL in an ImageQuant LAS 4000mini (GE Healthcare, Chicago, IL USA) as above.

### Cell envelope labeling

Chemical probes used in this work include alkDADA (Liechti et al., 2014), N-AlkTMM and O-AlkTMM (Foley et al., 2016) and FM4-64 FX (Invitrogen, Carlsbad, CA USA). AlkDADA was synthesized by the Einstein Chemical Biology Core and the TMM probes were kind gifts of Dr. Ben Swarts. Unless otherwise indicated, mid-log *M. smegmatis* or, where applicable, *B. subtilis, S. aureus, E. coli, L. lactis* or *C. crescentus* were labeled with 2 mM alkDADA, 250 μM N-AlkTMM, 50 μM O-AlkTMM or 5 µg/mL FM4-64FX for 15 min or 5 min in the case of *B. subtilis*. Unless otherwise indicated and where applicable, the bacteria were pre-incubated in the presence or absence of freshly-prepared chemicals (antibiotics or benzyl alcohol) before being subjected to the probes (see Table S1 and S2). Cells were pelleted by centrifugation, washed in PBST containing 0.01% BSA (PBSTB) and fixed for 10 min in 2% formaldehyde at room temperature. For alkDADA and O-AlkTMM, cells were further washed twice with PBSTB and subjected to CuAAC as described (García-Heredia et al., 2018). Unless otherwise specified, picolyl azide-Cy3 was used in Fig. 2D; picolyl azide carboxyrhodamine 110 was used in Figs. 3B, 3D and Figure 2–figure supplement 2, Figure 3–figure supplement 1, and Figure 3–figure supplement 3. Bacteria were then washed twice in PBSTB, once in PBST, and imaged (described below) or subjected to flow cytometry (BD DUAL LSRFortessa, UMass Amherst Flow Cytometry Core).

### Microscopy and image analysis

Bacteria were imaged on agar pads by either conventional fluorescence microscopy (Nikon Eclipse E600, Nikon Eclipse Ti or Zeiss Axioscope A1 with 100x objectives) or structured illumination microscopy (Nikon SIM-E/A1R with SR Apo TIRF 100x objective, UMass Amherst Light Microscopy Core).

To obtain the fluorescence intensity plots, the subcellular distribution of fluorescence was quantitated from images obtained by conventional fluorescence microscopy. The images were processed using Fiji and Oufti (Paintdakhi et al., 2016; Schindelin et al., 2012) as described (García-Heredia et al., 2018). The signal was normalized to length and total fluorescence intensity of the cell. Cells were oriented such that the brighter pole is on the right-hand of the graph. The intensity plots from Fig. 2A were made from 42<n<56 cells; for Fig. 3B, from 14<n<70 cells.

### Evaluation of membrane-bound peptidoglycan precursors

Wild-type or MurJ-depleted *M. smegmatis* (Gee et al., 2012) were grown to mid-log phase and membrane fractions were isolated as above. Precursors were extracted from each membrane fraction similar to previous publications (García-Heredia et al., 2018; Qiao et al., 2014; Qiao et al., 2017). Briefly, glacial acetic acid was added to 500 µL of fractionated lysate to a final volume of 1%. The sample was then transferred into a vial containing 500 µL of chloroform and 1 mL of methanol and left at room temperature for 1-2 hours with occasional vortexing. The mixture was centrifuged at 21,000x *g* for 10 min and the supernatant was transferred into a vial containing 500 µL of 1% glacial acetic acid (in water) and 500 µL chloroform, and vortexed for 1 min. The mixture was separated by centrifugation (900x *g* for 1 min at room temperature) and the organic phase was collected. Where applicable, the white interface was reextracted to recover additional organic material. The organic phase was dried under a nitrogen stream and the dried lipids were resuspended in 12 µL of DMSO. D-amino acid-containing, lipid-linked peptidoglycan precursors were biotinylated by subjecting organic extracts to an *in vitro* PBP4-mediated exchange reaction with biotin-D-lysine (Chem-Impex International, Wood Dale, IL USA; reagent was deprotected first) as described (Qiao et al., 2014). The products were separated by SDS-PAGE on an 18%, polyacrylamide gels then transferred to a PVDF membrane, blotted with streptavidin-HRP (diluted 1:10,000; Thermo Fisher Scientific, Waltham, MA USA) and visualized by ECL as above.

### Membrane glycolipid analysis

Sucrose density gradient fractions from wild-type *M. smegmatis* +/- 1 hour of 100 mM benzyl alcohol treatment or DivIVA-eGFP-ID with DivIVA depleted or not (Meniche et al., 2014) were subjected to lipid purification and analysis as previously described (Morita et al., 2005).

### Phylogenetic analysis

The phylogenetic tree was made in Adobe Illustrator (version 23.0.3) based on a phylogenetic tree generated with Mega (version 7.0.26; Kumar et al., 2018). Briefly, 16S rDNA sequences were obtained from NCBI (see Table S2) and aligned using ClustalW. The phylogenetic tree was generated using the Timura-Ney model with Gamma distribution and Bootstrap method (1000 replications). The taxonomic information was verified with the Interagency Taxonomic Information System (available online https://www.itis.gov/).

## Figure supplements

**Figure 1–figure supplement 1.**
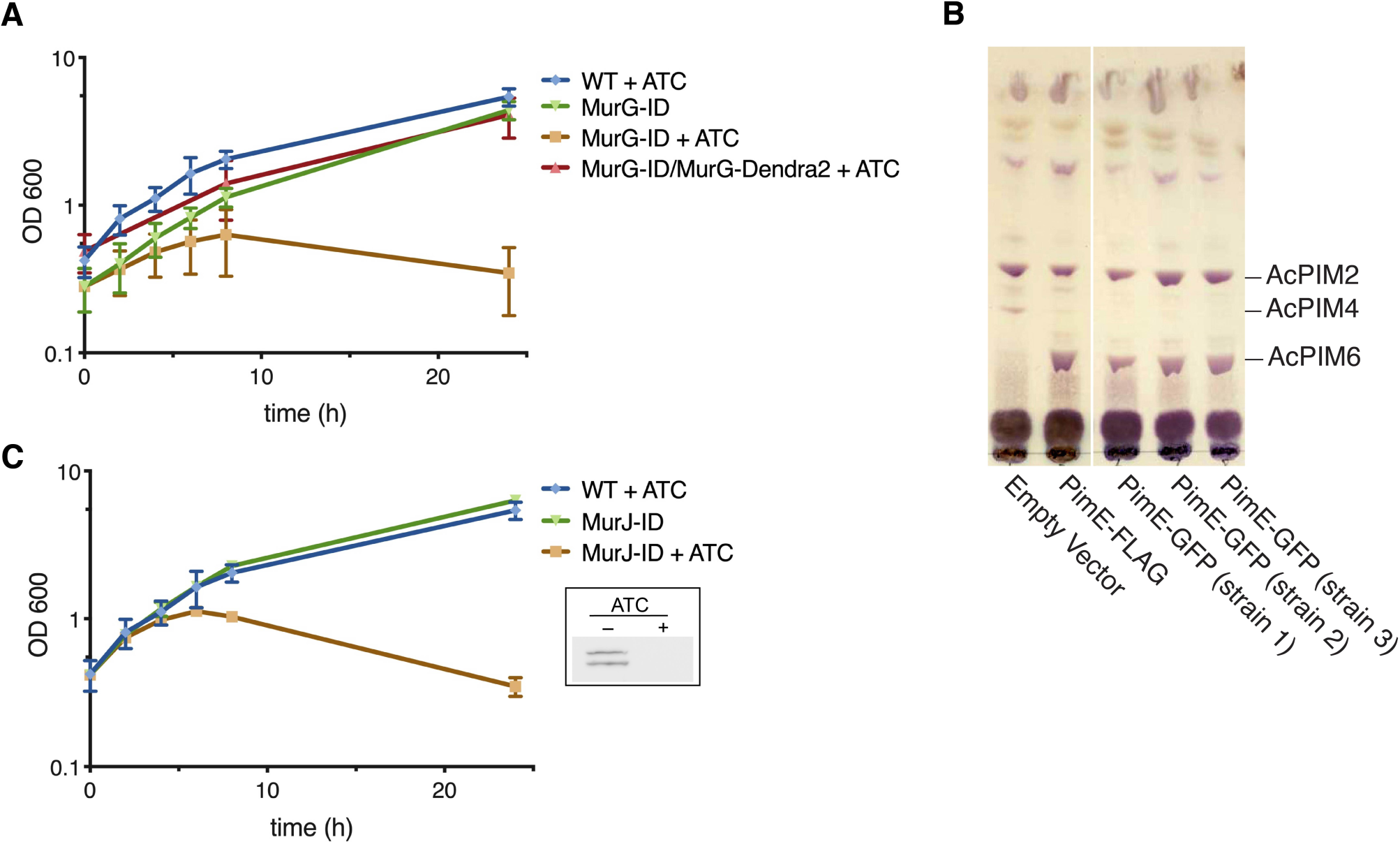
MurG-Dendra2 and PimE-GFP are functional, and MurJ can be depleted. (A) MurG-Dendra2 expression rescues depletion of endogenous protein. MurG depletion strain (Meniche et al., 2014) was transformed with a plasmid containing *murG-dendra2* and grown +/- 50 ng/mL anhydrotetracycline (ATC) to induce MurG-ID degradation. (B) PimE-GFP is functional. PimE is a mannosyltransferase involved in phosphatidylinositol mannoside (PIM) biosynthesis, converting AcPIM4 to more polar PIMs. *ΔpimE* was complemented with the indicated expression vectors. Glycolipids were extracted, purified, and separated by thin-layer chromatography. PIMs were visualized by orcinol staining. Similar to PimE-FLAG (Morita et al., 2006), PimE-GFP can restore the production of AcPIM6. (C) MurJ is depleted upon treatment with anhydrotetracycline (ATC). Depletion strain (MurJ-ID; carrying FLAG tag) was grown +/- ATC. Insert, immunoblot of lysates showing that MurJ is degraded after 8 hours of ATC treatment as reported (Gee et al., 2012).

**Figure 1–figure supplement 2.**
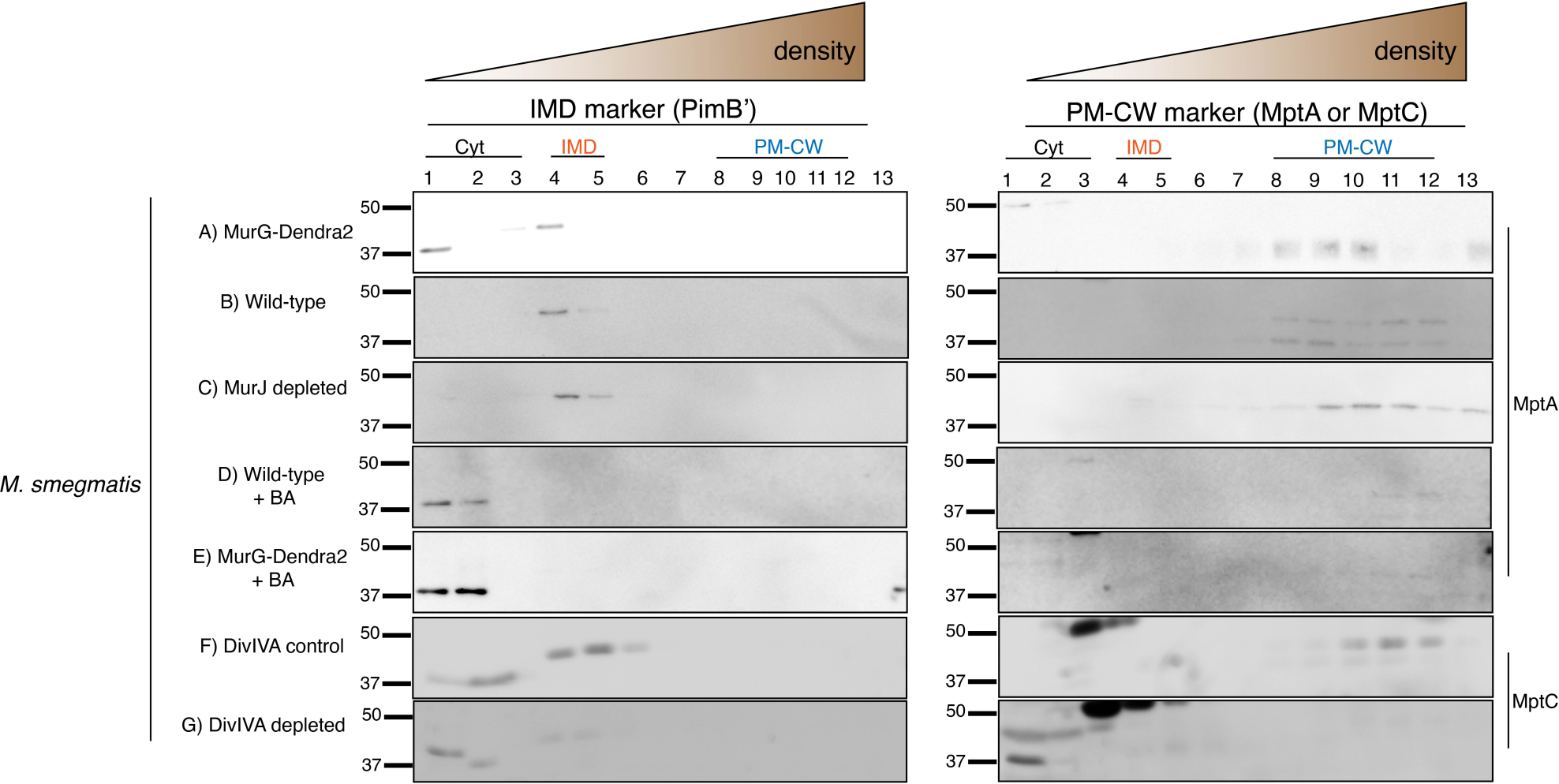
Western blot analysis of the IMD and the PM-CW membrane fractions separated by sucrose density sedimentation. Antibodies against PimB’ and MptA or MptC were used to detect the IMD and PM-CW, respectively (Hayashi et al., 2016; Sena et al., 2010). Analyzed fractions shown here are the same fractions of: A) MurG-Dendra2-expressing cells as shown in Fig. 1C; B) wild-type cells as shown in Fig. 1C and Fig. 2C; C) MurJ-depleted cells as shown in Fig. 2C; D) wild-type cells treated with benzyl alcohol (BA) as shown in Fig. 1C; E) MurG-Dendra2-expressing cells treated with benzyl alcohol as shown in Fig. 1C; F) and G) are DivIVA-eGFP-ID where DivIVA was depleted or not depleted as in Fig 4A. Samples from F) and G) were concentrated 10-fold by precipitating proteins in chloroform and water. PimB’ appears as ∼45 kDa band in the IMD fractions but disappears upon BA treatment or DivIVA depletion. We do not yet know the reason for this disappearance. The band at ∼37 kDa, which is visible especially in the cytoplasmic fractions of BA-treated cells is a non-specific protein (Hayashi et al., 2016). MptA and MptC appear as ∼42 kDa band in the PM-CW fractions, which became fainter after BA treatment or depletion of DivIVA.

**Figure 1–figure supplement 3.**
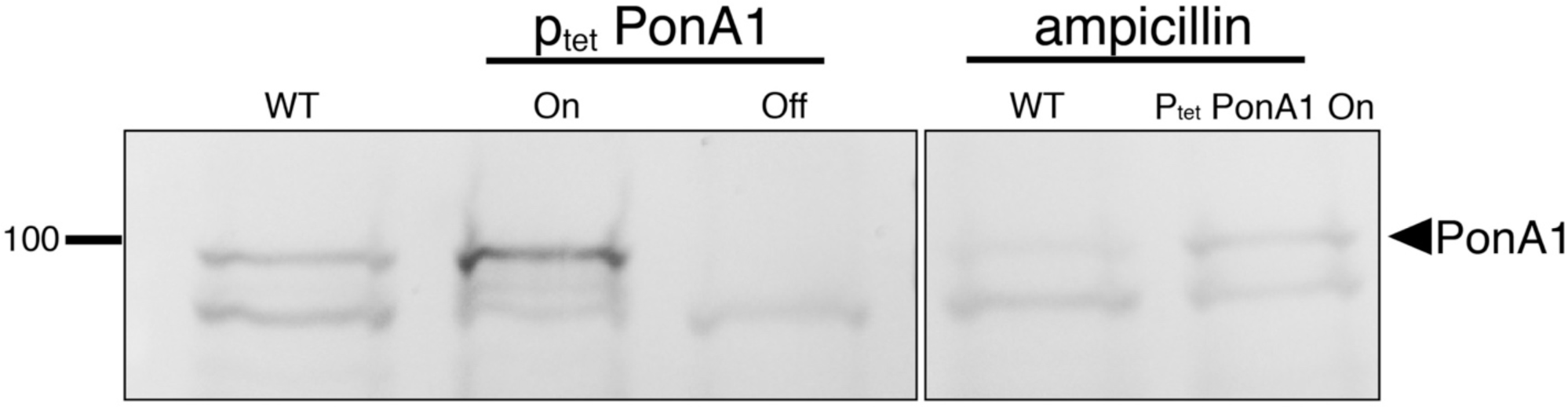
Visualization of PonA1 using Bocillin-FL. To assess specificity of the fluorescent signal, membrane fractions were pre-incubated +/- 16 µg/mL ampicillin and 5 µg/mL of the β-lactamase inhibitor clavulanate (García-Heredia et al., 2018) for 30 min at room temperature. 100 µg of total membrane proteins were incubated with 40 µM of Bocillin-FL for 30 min, then separated by SDS-PAGE (12% gel) and visualized by in-gel fluorescence to detect PBPs. WT, wild-type *M. smegmatis*; p_tet_*ponA1* expressed (On) or depleted (Off) with addition of anhydrotetracycline (ATC) as described (Hett et al., 2010).

**Figure 2–figure supplement 1.**
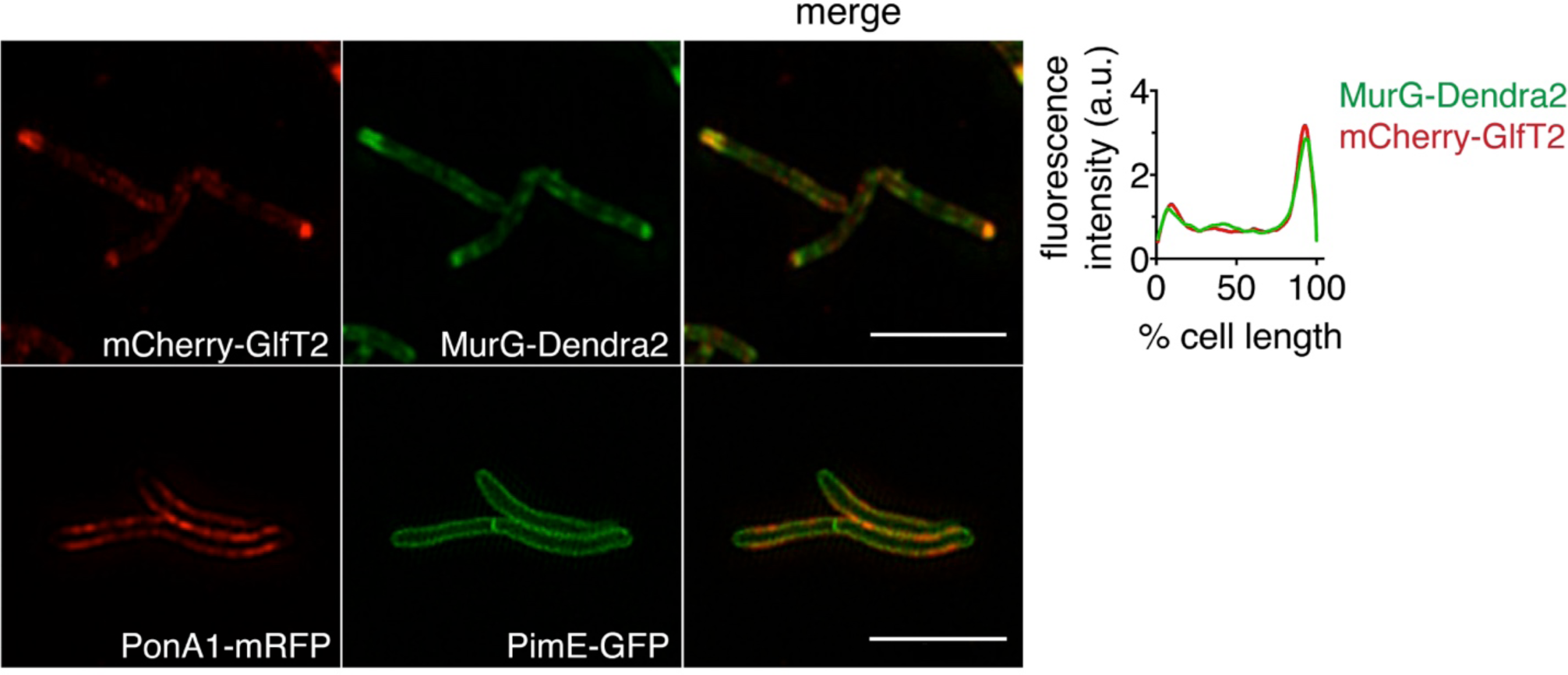
MurG-Dendra2 and PonA1-mRFP colocalize with IMD and PM-CW reporters, respectively. *M. smegmatis* coexpressing MurG-Dendra2 and mCherry-GlfT2 or PonA1-mRFP and PimE-GFP were imaged by SIM-E. Scale bars, 5 µm. Graph, the fluorescence intensity profiles of the proteins were quantitated as in Fig 2A from images captured with conventional microscopy.

**Figure 2–figure supplement 2.**
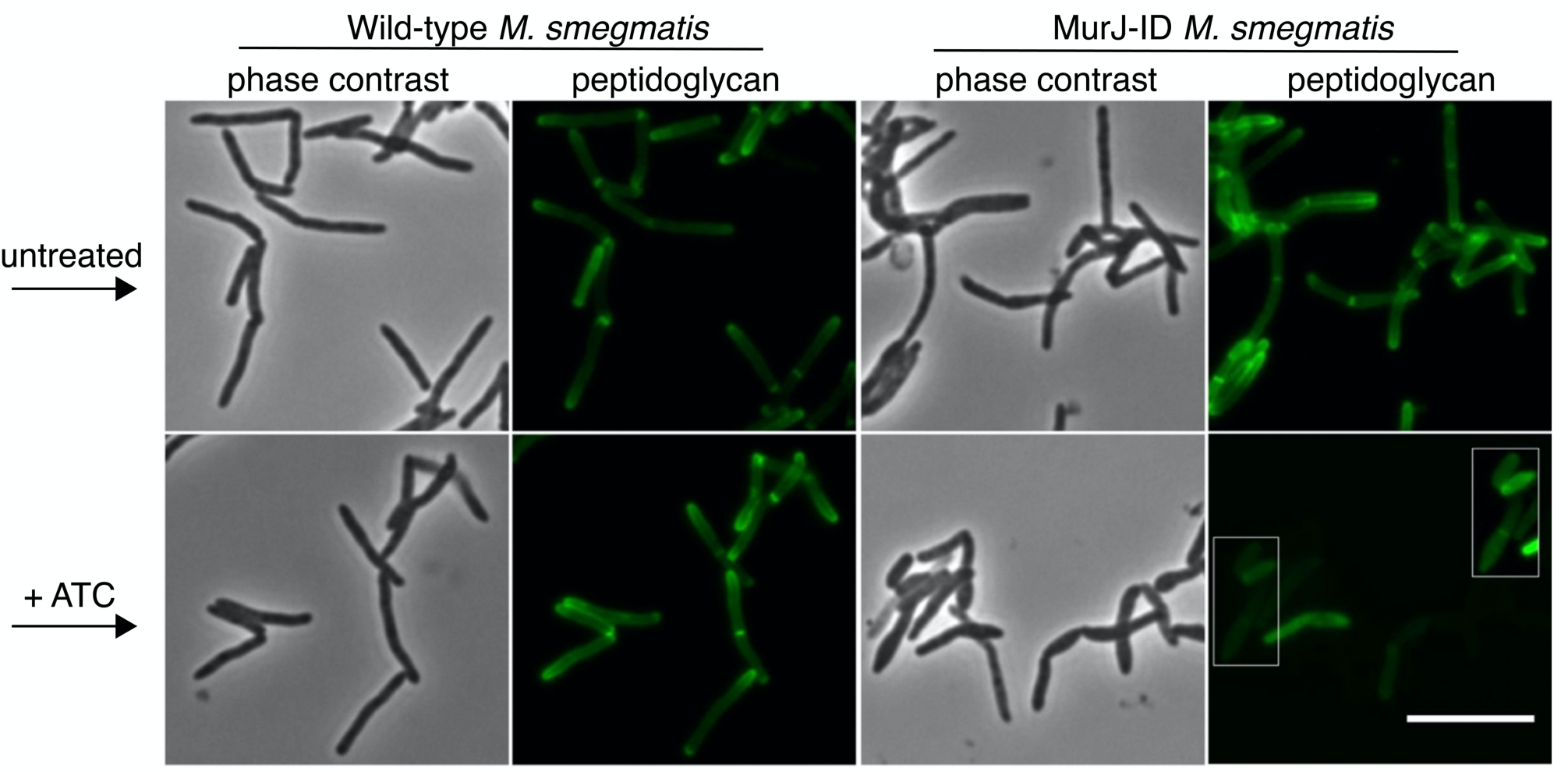
Depletion of MurJ alters amount and location of nascent peptidoglycan. Wild-type (left) or MurJ-ID (depletion strain; right) *M. smegmatis* in early log phase were treated +/ anhydrotetracycline (ATC) then incubated with alkDADA. Bacteria were washed, fixed, subjected to CuAAC and imaged by conventional fluorescence microscopy. Scale bar, 5 µm.

**Figure 3–figure supplement 1.**
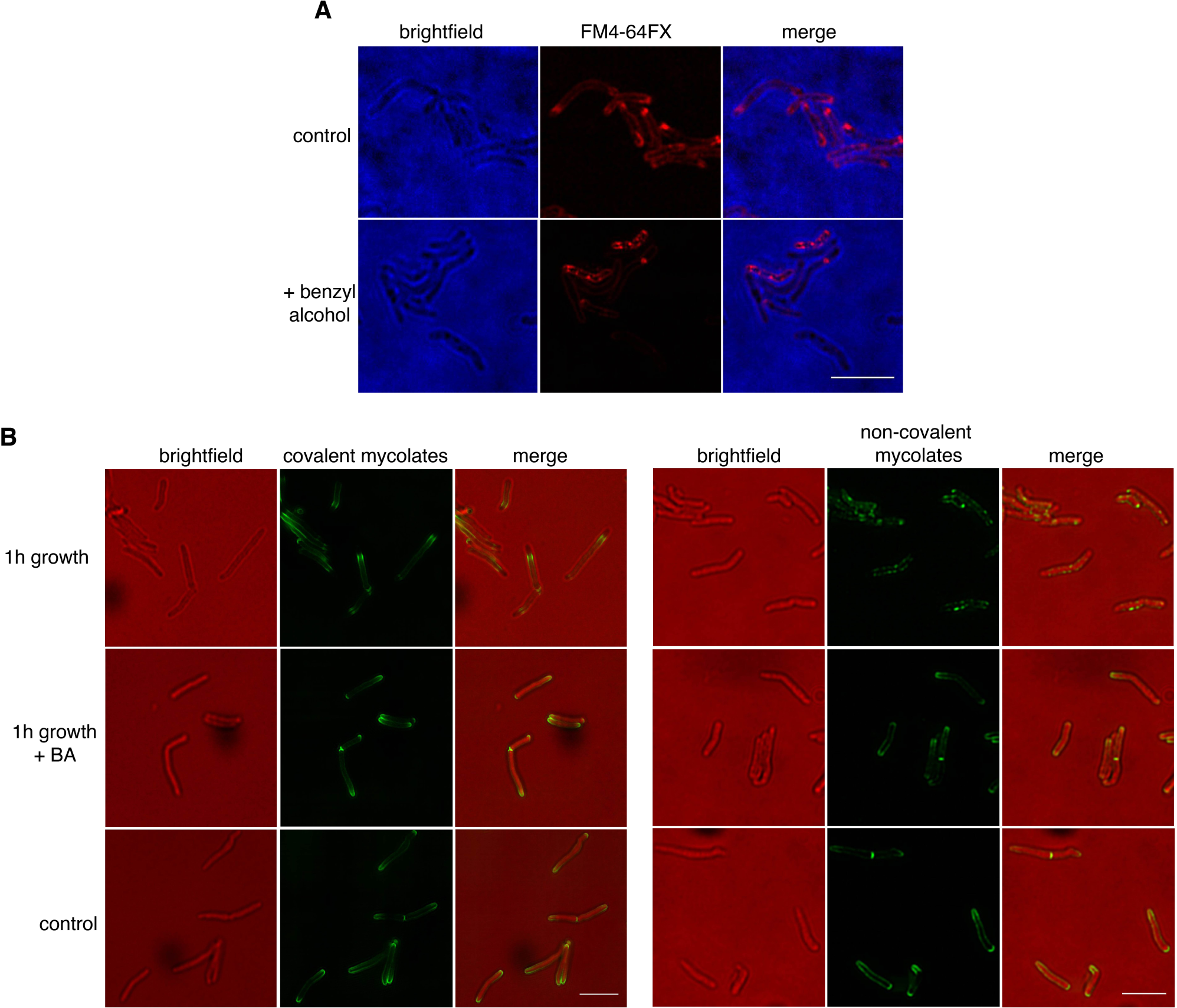
(A) Benzyl alcohol alters FM4-64FX distribution. Wild-type *M. smegmatis* was labeled with FM4-64FX, then washed and incubated with benzyl alcohol. Cells were then imaged with SIM-E. (B) Benzyl alcohol halts cell elongation but does not otherwise impact the localization of mycomembrane probes. Wild-type *M. smegmatis* was incubated with O-AlkTMM (left) or N-AlkTMM (right) to label covalent or noncovalent mycolates, respectively, then washed and subjected or not to benzyl alcohol for 1 hour. Bacteria were washed, fixed, subjected to CuAAC and imaged by conventional fluorescence microscopy. As a control, bacteria were not treated with benzyl alcohol for 1 hour, *i.e.* fixed immediately after probe incubation (bottom panel). All scale bars, 5 µm.

**Figure 3–figure supplement 2.**
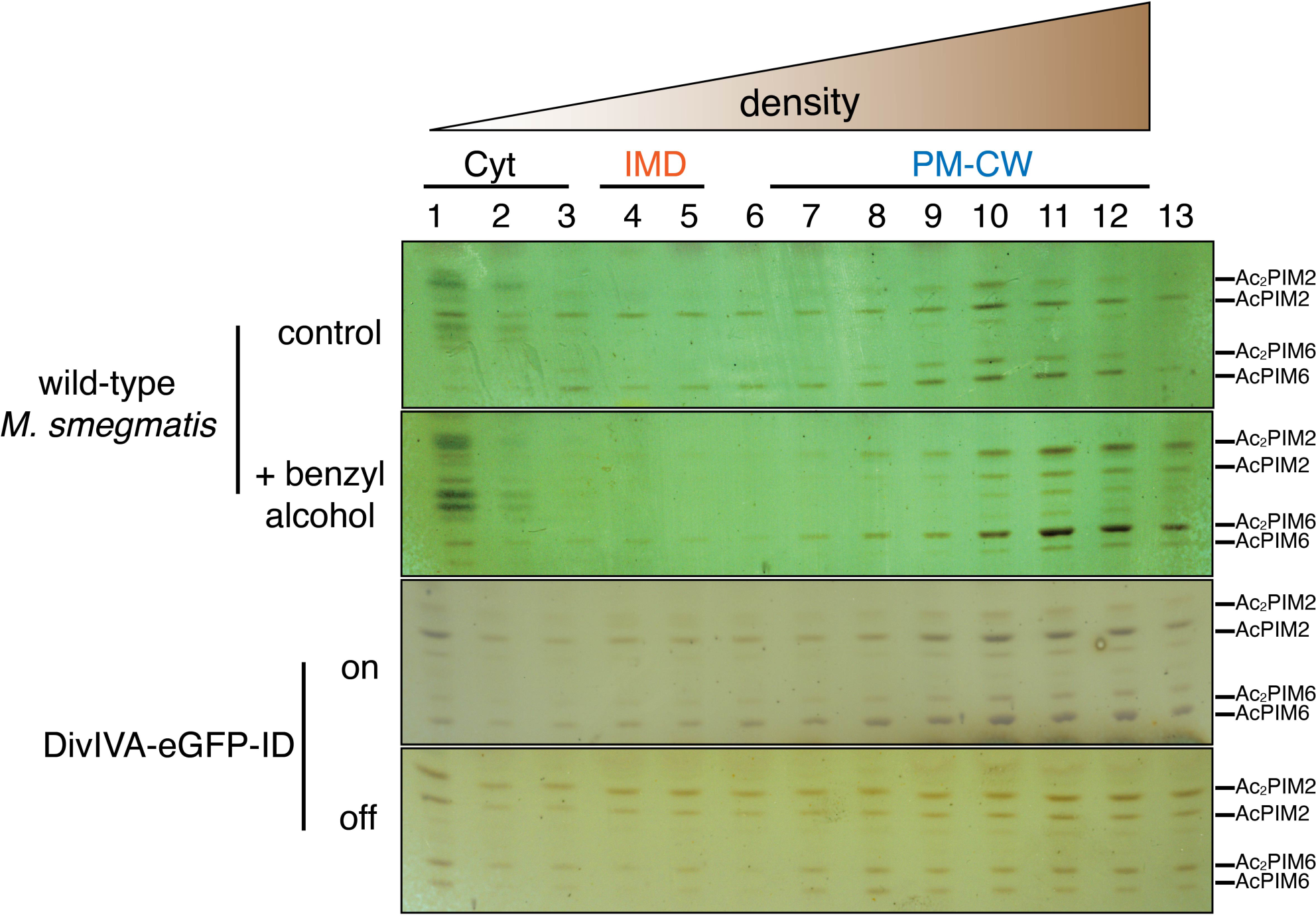
Benzyl alcohol and depletion of DivIVA affect the distribution of membrane glycolipids. Wild-type or DivIVA-eGFP-ID *M. smegmatis* were treated +/- benzyl alcohol or ATC, respectively. PIMs were visualized by orcinol staining as in Figure 1. –figure supplement 1. AcPIM2 and AcPIM6 present during normal growth in the IMD and are diminished upon benzyl alcohol treatment. Depletion of DivIVA is accompanied by an enrichment of PIM2 species and depletion of PIM6 species. This change in lipids is currently under investigation.

**Figure 3–figure supplement 3.**
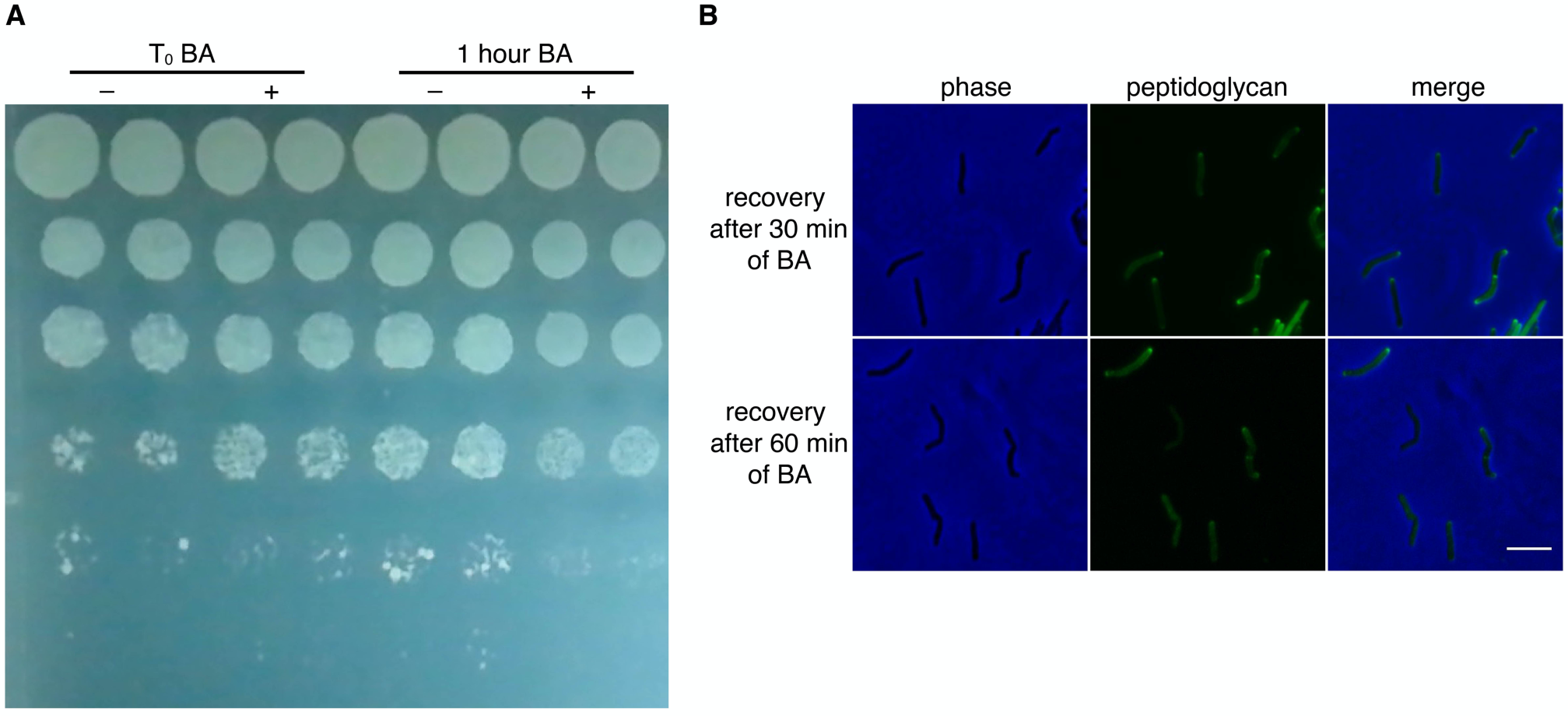
*M. smegmatis* survives benzyl alcohol (BA) treatment. (A) *M. smegmatis* was treated with benzyl alcohol then washed in PBS. Ten-fold serial dilutions were spotted on LB agar. Shown are cells immediately after addition of benzyl alcohol (T_0_) and after 1 hour of exposure. (B) *M. smegmatis* were treated with benzyl alcohol for 30 or 60 min, washed and incubated for 2 hours in fresh 7H9. Cells were then labeled with alkDADA for 15 min and then fixed and alkynes were detected by CuAAC to reveal active peptidoglycan metabolism. Scale bar 5 µm.

**Figure 3–figure supplement 4.**
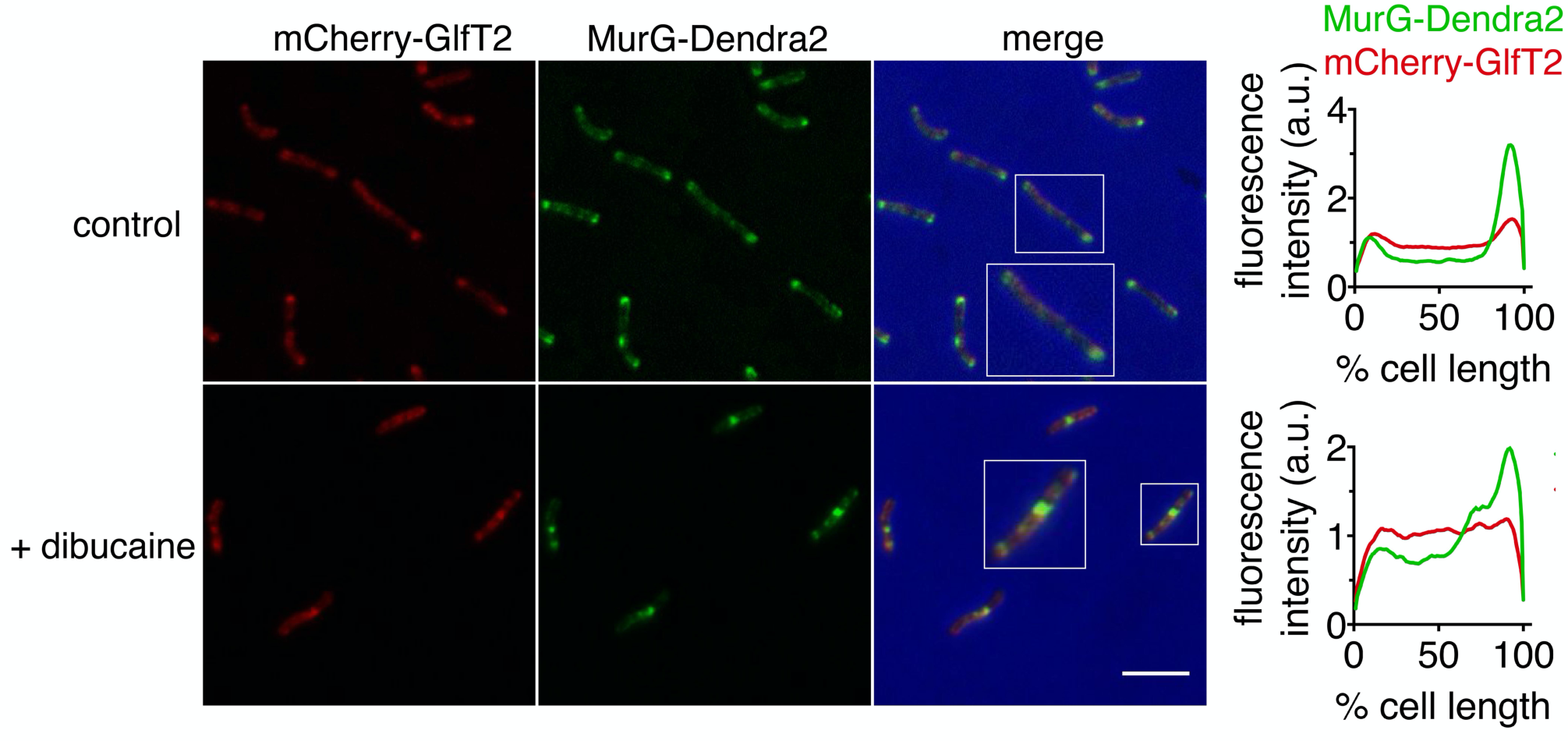
Dibucaine impacts the localization of MurG and GlfT2. *M. smegmatis* coexpressing the IMD-associated MurG-Dendra2 and mCherry-GlfT2 was treated or not with dibucaine. Images from conventional microscopy show these proteins lose their polar enrichment when treated with dibucaine. Scale bar, 5 µm. Right, the fluorescence intensity profiles of the fusion proteins were determined as in Figure 2A.

**Figure 4–figure supplement 1.**
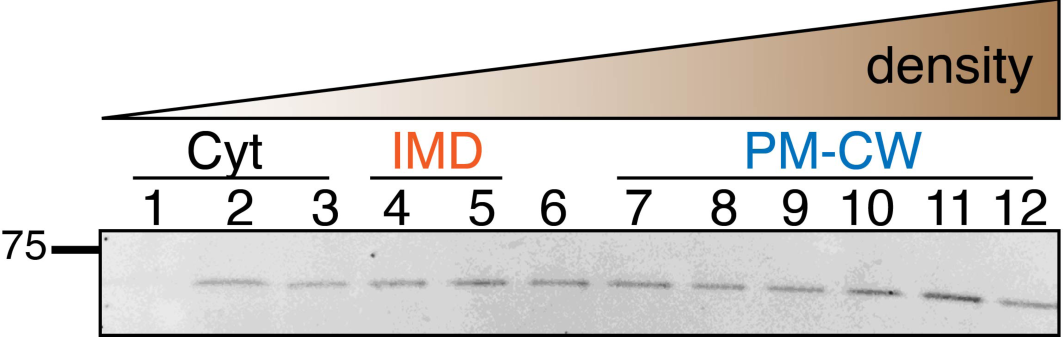
Cell wall is required for partitioning of MurG. *M. smegmatis* expressing MurG-Dendra2 were spheroplasted as in Fig. 4A and then lysed by nitrogen cavitation. The crude lysates were separated in a density gradient and blotted in a polyacrylamide gel. MurG-Dendra2 distributes widely along the membrane fractions consistent with the broad distribution of cellular material across the density gradient (see Fig. 4A), which contrasts with its normal partitioning (see Fig. 1C).

## Supplementary information

**Table S1.**
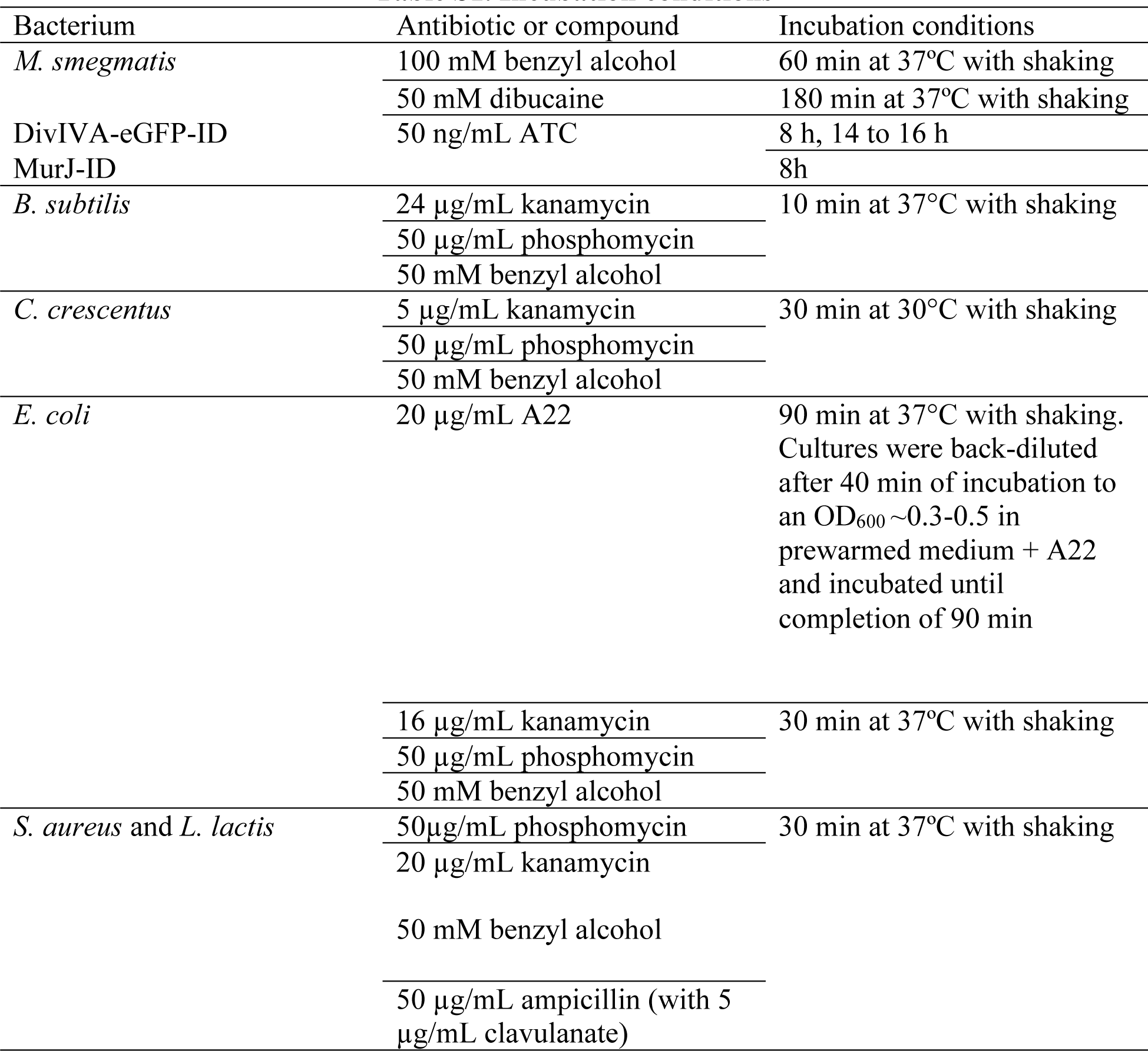
Incubation conditions.

**Table S2.** Incubation conditions of probes to metabolically label the cell envelope.

| Bacterium | Probe | Labeling conditions |
| --- | --- | --- |
| *M. smegmatis* | alkDADA<br>N-Alk-TMM<br>O-Alk-TMM | Unless otherwise indicated,<br>15 min with shacking at 37°C |
| *E. coli*<br>*B. subtilis*<br>*S. aureus*<br>*L. lactis* | alkDADA | Unless otherwise indicated, 5<br>min with shacking at 37°C |
| *C. crescentus* |  | Unless otherwise indicated, 5<br>min with shacking at 30°C |

**Table S3.** NCBI Accession numbers from 16S rDNA.

| Bacterium | NCBI accession number |
| --- | --- |
| *M. smegmatis* | AB305022.1 |
| *B. subtilis* | NR_112116.2 |
| *S. aureus* | NR_118997.2 |
| *L. lactis* | FJ348447.1 |
| *E. coli* | NR_024570.1 |
| *C. crescentus* | NR_037099.1 |

**Table S4.** Key Resources.

| Reagent type | Designation | Source or reference | Additional information |
| --- | --- | --- | --- |
| Strain (*M. smegmatis* mc<sup>2</sup>155) | *M. smegmatis* | NC_008596 in GenBank | Wild-type *M. smegmatis* |
| Strain (*M. smegmatis*) | MurG-Dendra2 | This study | The mutant was generated as described in supplementary material and methods. |
| Strain (*M. smegmatis*) | mCherry-GlfT2 | (Hayashi et al., 2016) | See reference for details. |
| Strain (*M. smegmatis*) | PonA1-mRFP | (Kieser et al., 2015) | Obtained from Dr. Eric Rubin (Harvard SPH) and Dr. Hesper Rego (Yale Med) |
| Strain (*M. smegmatis*) | PimE-GFP | This study | The mutant was generated as described in supplementary material and methods. |
| Strain (*M. smegmatis*) | MurG-ID depletion strain | (Meniche et al., 2014) | Obtained from Dr. Chris Sassetti (U Mass Med) |
| Strain (*M. smegmatis*) | MurJ-ID (MviN) depletion strain | (Gee et al., 2012) | Obtained from Dr. Chris Sassetti (U Mass Med) |
| Strain (*M. smegmatis*) | p<sub>tet</sub>*ponA1* | (Hett et al., 2010) | Obtained from Dr. Eric Rubin (Harvard SPH) |
| Strain (*M. smegmatis*) | DivIVA-eGFP-ID | (Meniche et al., 2014) | Obtained from Dr. Chris Sassetti (U Mass Med) |
| Strain (*M. smegmatis*) | mCherry-GlfT2/DivIVA-eGFP-ID | This study |  |
| Strain (*B. subtilis* JH642) | *B. subtilis* | NZ_CP007800 in GeneBank |  |
| Strain (*C. crescentus*) | *C. crescentus* | NA 1000 | Obtained from Dr. Peter Chien (U Mass Amherst) |
| Strain (*E. coli* K12) | *E. coli* K12 | MG1655 |  |
| Strain (*S. aureus*) | *S. aureus* | ATCC BA-1718 | Obtained from Dr. Thai Thayumanavan (U Mass Amherst) |
| Strain (*L. lactis*) | *L. lactis lactis* | NRRL B633 |  |
| Chemical compound | alkyne-D-alanine-D-alanine (alkDADA or EDA-DA) | (Liechti et al., 2014) | Synthesized by the Chemical Synthesis Core Facility at Albert Einstein College of Medicine (NY, USA) following the referenced protocols |
| Chemical compound | O-alkyne-trehalose monomycolate (O-AlkTMM) | (Foley et al., 2016) | Obtained from Dr. Benjamin Swarts (Central Michigan University) |
| Chemical compound | N-alkyne-trehalose monomycolate (N-AlkTMM) | (Foley et al., 2016) | Obtained from Dr. Benjamin Swarts (Central Michigan University) |
| Software, algorithm | MATLAB codes | (García-Heredia et al., 2018) | Scripts designed for MATLAB to analyze the fluorescence profiles along a cell body from data collected in Oufiti<sup>28</sup>. |
| Chemical compound | Fmoc-D-Lys(biotinyl)-OH BDL precursor | Chem-Impex International (Wood Dale, IL USA) | Deprotected as described in<sup>13</sup> to yield BDL |
| Chemical compound | A22 (S-3,4-Dichlorobenzylisothioureahydrochloride) | Sigma-Aldrich, St. Louis, MO USA | Dissolved in water and kept at -20°C |
| Recombinant DNA reagent | PBP4 plasmid | (Qiao et al., 2014) | Obtained from Dr. Suzanne Walker (Harvard Med) |

